# Automated high-throughput profiling of single-cell total transcriptome with scComplete-seq

**DOI:** 10.1101/2024.03.12.584729

**Authors:** Fatma Betül Dinçaslan, Shaun Wei Yang Ngang, Rui Zhen Tan, Lih Feng Cheow

## Abstract

Detecting the complete portrait of the transcriptome is essential to understanding the roles of both polyadenylated and non-polyadenylated RNA species. However, current efforts to investigate the heterogeneity of the total cellular transcriptome in single cells are limited by the lack of an automated, high-throughput assay that can be carried out on existing platforms. To address this issue, we developed scComplete-seq, a method that can easily augment existing high-throughput droplet-based single-cell mRNA sequencing to provide additional information on the non-polyadenylated transcriptome. Using scComplete-seq, we have successfully detected long and short non-polyadenylated RNAs at single-cell resolution, including cell-cycle-specific histone RNAs, cell-type-specific short non-coding RNA, as well as enhancer RNAs in cancer cells and PBMCs. By applying scComplete-seq, we have identified changes in both coding and non-coding transcriptome in PBMCs during different stimulations. Measuring the enhancer RNA expression also revealed the activation of specific biological processes and the transcription factors regulating such changes.

## INTRODUCTION

One of the greatest technological innovations that have advanced our understanding of cell biology in the past decade is the development of high-throughput single-cell mRNA-sequencing (scmRNA-seq) platforms. Today, commercial scmRNA-seq platforms (e.g. 10X Genomics Chromium) can routinely profile the coding transcriptome of ∼10,000 single cells. The convenient workflow, quick turnaround, and affordable cost have led to very wide adoption among the scientific communities for interrogating cellular heterogeneities in various applications, and have in turn spurned international efforts such as the human cell atlas program.

A current limitation of most current high-throughput scmRNA-seq platforms is that they only allow the profiling of polyadenylated RNA species, due to the common use of oligo(dT) primers for reverse transcription. However, the polyadenylated RNA species (which include mRNA and a subset of long non-coding RNA (lncRNA)) only account for 3-7% of the total transcriptome^1^. There is an abundance of other non-polyadenylated RNA species including histone RNA, ribosomal RNA (rRNA), transfer RNA (tRNA), small nuclear RNA (snRNA), and microRNA (miRNA), which play important functional and regulatory roles in cells. To fully profile the transcriptomic heterogeneity among cells, and to understand the roles of these non-coding RNA in cellular diversity, there is a pressing need for technologies that can simultaneously profile both polyadenylated and non-polyadenylated RNA species at the single-cell level, in a highly automated, accessible, and high-throughput manner.

There have been several attempts to develop single-cell non-polyadenylated RNA sequencing capabilities in recent years. Small-seq is a ligation-based approach that enables the sequencing of small RNA in single cells^2^. Using a half-cell genomics approach, Wang *et. al*. combined Small-seq with scmRNA-seq to perform paired analysis of microRNA and mRNA from single cells^3^. As the ligation-based approach for sequencing small RNA involved a large number of steps, they could only be performed in microwell plates, which increased the cost-per-cell and limited the throughput to 100s of single cells. On the other hand, Smart-seq Total^4^ and VASA-seq^5^ made use of Poly(A) Polymerase (PAP) enzymes to polyadenylate all RNA before reverse transcription with oligo(dT) primers to capture all RNA species. Multiple reagent addition steps were still needed in the Smart-seq Total and VASA-seq approach. This was achieved by manual sequential addition of reagents into microwell plates in Smart-seq Total^4^ and utilization of three custom microfluidic chips and custom barcoded polyacrylamide beads for sequential reagent addition into cell-containing droplets in VASA-drop^5^. To date, there is no automated high-throughput method that is generally available to the broader scientific community for profiling the total transcriptome of single cells.

In this manuscript, we demonstrate a novel method to enable total RNA sequencing that is compatible with a commercially available high-throughput single-cell analysis platform (10X Genomics Chromium). The key innovation is the incorporation of PAP enzyme and use of locked-nucleic-acid modified template-switching-oligos (LNA-TSO) to enable single-step cell-lysis, *in vitro* RNA polyadenylation, reverse transcription, and template-switching reaction in droplets. We demonstrate the efficient recovery of non-coding RNA that characterized the cell types and cell cycle with this single-cell approach. Applying this method to peripheral blood mononuclear cells (PBMC), we discovered unique patterns of tRNA and snRNA associated with cellular subtypes and their dynamic changes in response to different stimuli. Moreover, enhancer RNAs could be simultaneously profiled with mRNA in this method, providing important insights into the transcriptional regulation by enhancers. Single-cell Complete RNA sequencing (scComplete-seq) is a versatile technology that can readily enable the exploration of the comprehensive transcriptome at single-cell resolution for various applications.

## RESULTS

Droplet microfluidics are very attractive high-throughput platforms for scmRNA-seq due to the ease of automatically generating hundreds of thousands of droplets containing cells and the significant reduction of reagent cost. A key to the commercial feasibility of droplet-based scmRNA-seq platforms is achieving a one-step process of in-droplet cell lysis and complete cDNA synthesis. This allows a very easy single-chip device operation that can be operated by even non-specialists.

Various single-cell total RNA sequencing technologies have been developed in recent years, but all of them require multiple sequential steps that pose considerable challenges to their miniaturization, automation, and scaling. If one could combine all the steps from single-cell lysis to total cDNA synthesis into a single reaction, a single-chip droplet-based single-cell comprehensive RNA sequencing platform could be realized.

The addition of PAP enzymes to polyadenylate the 3’ ends of RNA is an elegant approach that would enable all RNA to be reverse transcribed using oligo(dT) primers. Unlike non-treated cellular RNA, *E*.*Coli* PAP enzyme-treated RNA showed distinct bands at ∼1,900nt and ∼4,000nt when RNA was isolated with oligo(dT) magnetic beads (Fig. 1a). This demonstrates that RNA species that do not possess natural poly-A tails (e.g. 18S and 28S rRNA) could be interrogated through *in vitro* polyadenylation step.

**Figure 1:**
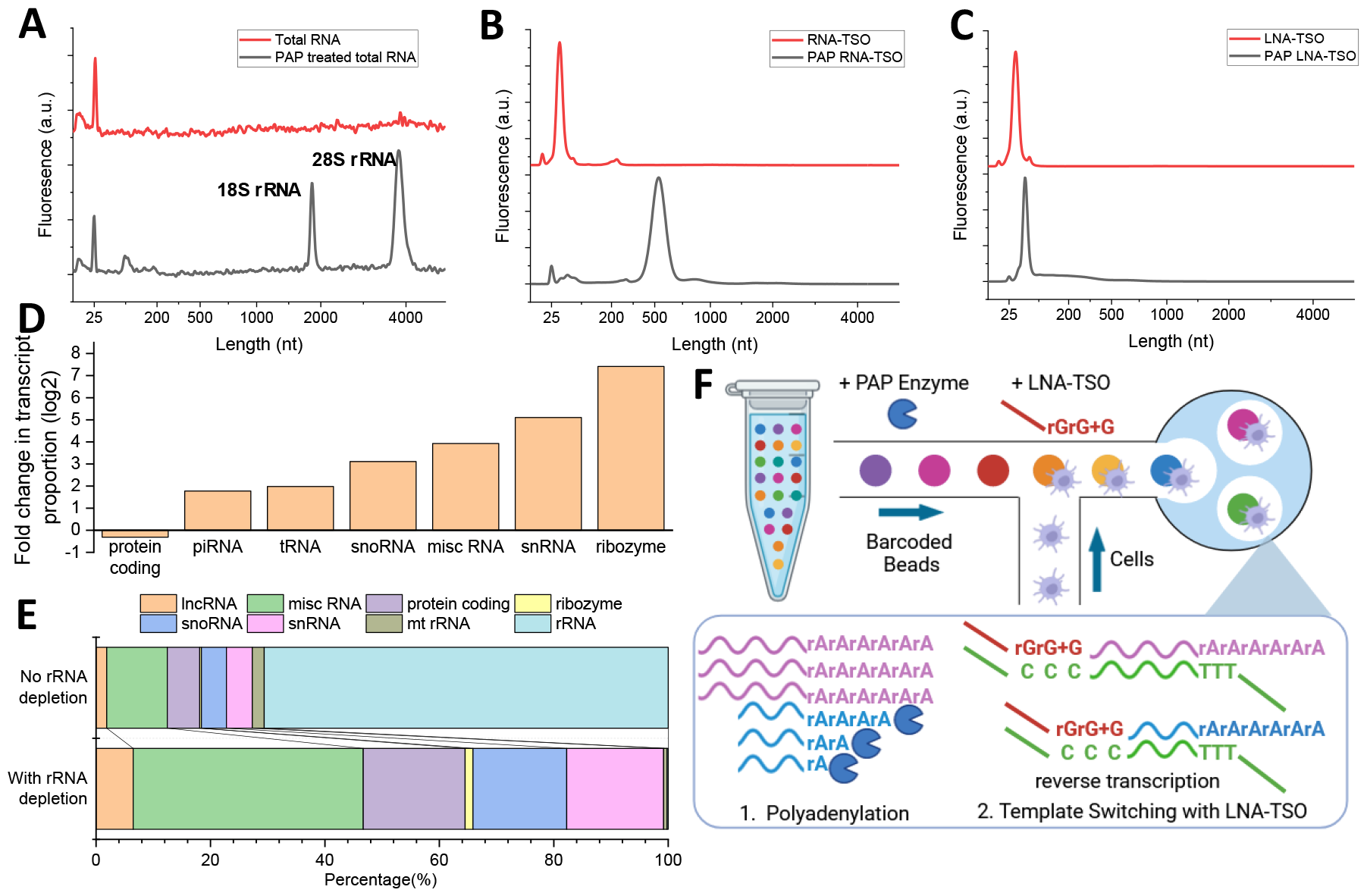
Key concepts for enabling scComplete-seq. **(A)** Treating total RNA with PAP enzyme allows polyadenylation of all RNA species and allows non-coding RNA including rRNA to be interrogated. **(B)** RNA-TSO is extended by PAP due to its 3’ ribonucleotide. (C) Polyadenylation of LNA-TSO is prevented by replacing its 3’ ribonucleotide with locked-nucleic acid. (D) Enrichment of non-polyadenylated RNA species in bulk Complete-seq. (E) Post-library rRNA depletion increases the sensitivity of detecting other RNA species in Complete-seq. (F) Schematic workflow of implementing scComplete-seq on an automated high-throughput single-cell platform.

The Switching Mechanism At the 5’ end of RNA Template (SMART)^6^ technology is the most widely-used strategy for generating amplification-ready cDNA. In SMART-seq, the poly(A) tail of RNA is primed using an oligo(dT) primer, reverse transcriptase with terminal transferase activity adds non-templated cytosines to the 3’ end of the cDNA, which allows template switching through annealing of a template-switching oligo with riboguanosine at its 3’ end (RNA-TSO). However, RNA-TSO is fundamentally incompatible with PAP as it is extended during co-incubation (Fig. 1b).

One strategy to prevent the inadvertent extension of TSO during the one-step reaction is to utilize TSO that does not end with ribonucleotides. Previous work^7^ has shown that TSO that contains LNA as the penultimate base in the 3’ end (riboguanosine as the third and second last base, LNA-guanosine as the last base) can facilitate template switching. We tested the incubation of LNA-TSO with PAP and confirmed the absence of extension (Fig. 1c).

Combining the above observations, we hypothesize that cell lysis, *in-vitro* RNA polyadenylation, reverse transcription, and template switching could be combined in a single reaction. We formulated a reaction mix where cell lysis and reverse transcription reagents were supplemented by PAP enzymes and LNA-TSO (Complete-seq workflow), followed by library preparation and next-generation sequencing. The result of bulk Complete-seq was compared with conventional mRNA-seq. We found that Complete-seq captured significantly more amounts of non-polyadenylated RNA of various biotypes (Fig. 1d), thus offering a simple yet powerful tool for comprehensive transcriptome profiling. Among the reads mapping to the transcriptome in Complete-seq, ∼70% mapped to rRNA, consistent with rRNA being the most abundant non-coding RNA (Fig. 1e). The large proportion of rDNA could mask the measurement of the other low abundant species. We applied a post-library depletion strategy (SEQuoia RiboDepletion, BioRad) to remove fragments derived from ribosomal RNA from the amplified library. This method reduced the cytoplasmic and mitochondria rRNA percentage from 70% and 2% to 0.2% and 0.5% respectively (Fig. 1e), and allowed considerably more protein-coding and non-coding RNA to be measured.

The ability to obtain cDNA from cells in a single step in Complete-seq is highly attractive for high throughput single-cell implementation. We implement this strategy on a commercial scmRNA-seq platform (10X Genomics Chromium). The original lysis-reverse transcription mix was supplemented with PAP enzyme and ATP, whereas the RNA-TSO was replaced with a custom LNA-TSO. The droplet incubation time was extended to facilitate the additional polyadenylation step. After in-droplet cDNA synthesis, the droplets were de-emulsified to recover cDNA, and cDNA preamplification was performed according to the 10X Genomics Chromium protocol (Fig. 1f). Due to the large sizes of mRNA, they were typically recovered from the large fractions (>600bp) in conventional scmRNA-seq. However, we expect the small fraction (<600bp) to contain abundant information about small non-coding RNA. Thus, both the large and small fractions were recovered and sequenced separately in scComplete-seq.

### Validation of scComplete-seq in cancer cell lines

To validate scComplete-seq, we performed experiments on three well-characterized cancer cell line populations. These consisted of a breast cancer cell line (MCF7), a myelogenous leukemia cell line (K562), and an embryonic kidney cell line (HEK293T). These cells were labeled with individual hashtags for identification and pooled for single-cell sequencing. We performed parallel scmRNA-seq and scComplete-seq experiments to benchmark the assay. Library preparation was performed on the large fraction of the scmRNA-seq samples while both the large and small fractions were sequenced in the scComplete-seq samples.

We first asked if scComplete-seq preserved the information contained within scmRNA-seq. Here, we focused on the large fraction of scComplete-seq as it was expected to contain the majority of the large mRNA species. Both scmRNA-seq and scComplete-seq detected a comparable number of unique mRNA per cell (scmRNAseq average: 1912 HEK293T, 1514 MCF7, 2467 K562, scComplete-seq average: 1853 HEK293T, 1430 MCF7, 2322 K562). Within each cell type, there was a strong correlation in mRNA gene expression between scmRNA-seq and scComplete-seq, indicating preservation of mRNA gene expression information in scComplete-seq (Fig 2a, Supp Fig. 1). However, at a whole-transcriptome level, we observed several species that were observed much more frequently in the scComplete-seq libraries. These included many histone genes (e.g. *H2BC11, H3C2, H2AC17, H4C8, H2BC18, H2AC11*), signal recognition particle RNA (e.g. *RN7SL1*) and various species mapping to tRNA (e.g. *tRNA-Gly-CCC-2-1, tRNA-Val-TAC-3-1*). tRNA and signal recognition particle RNA lack polyadenylation as they are transcribed by RNA polymerase III. Many histone genes were also known to lack polyadenylation. The observation of these non-polyadenylated RNA species in scComplete-seq libraries was strong evidence supporting the working principles of scComplete-seq.

**Figure 2.**
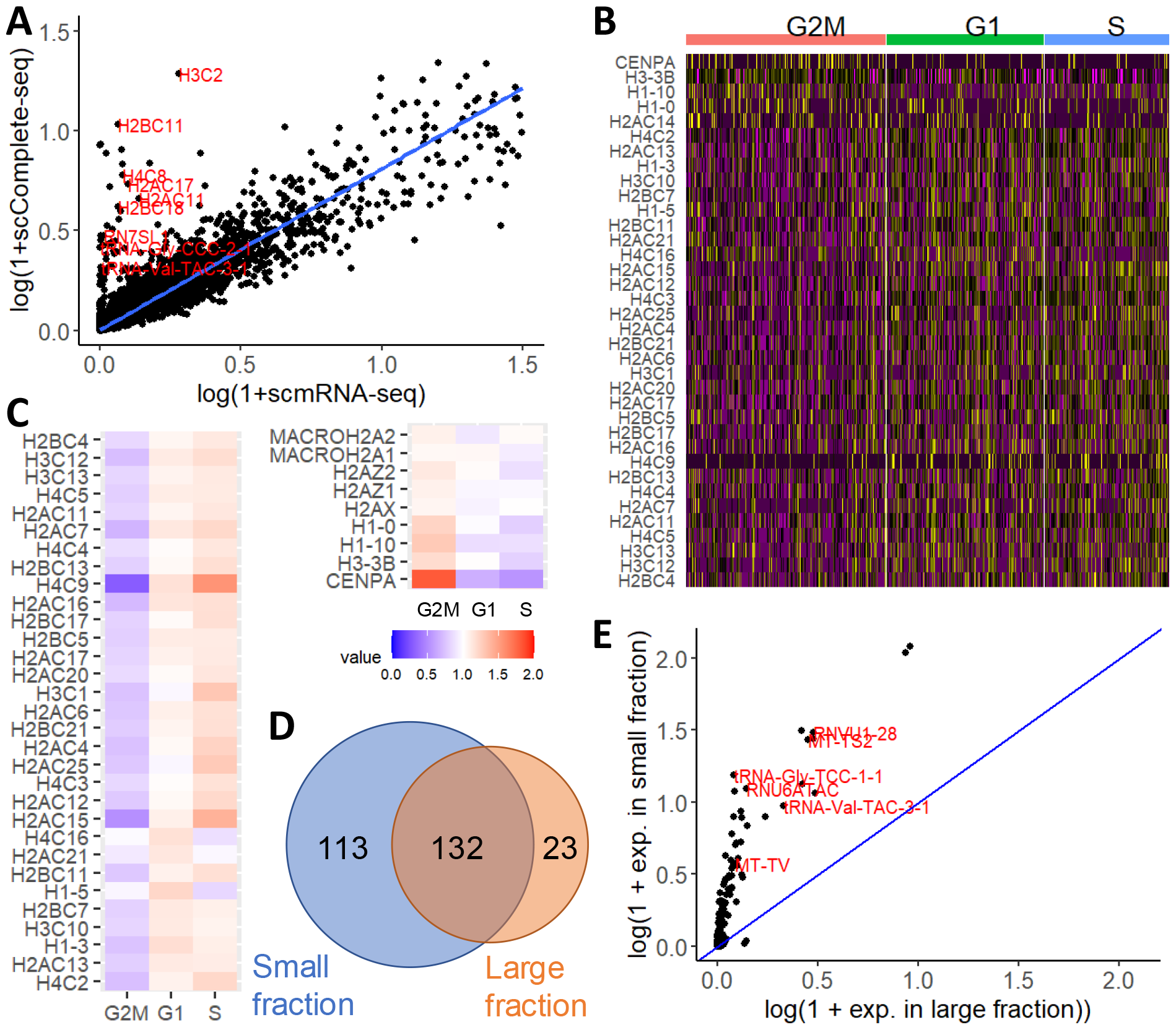
Detection of non-polyadenylated RNA in scComplete-seq. **(A)** Aggregated read counts obtained from scComplete-seq versus scmRNA-seq in HEK293T cells. **(B)** Histone gene expression for HEK293T, K562, and MCF7 cells in G2M, G1, and S phase. **(C)** Expression of non-polyadenylated (left) and polyadenylated histone RNA according to cell cycle phase. **(D)** Overlap between the number of short non-coding RNA detected in the small and large fraction. **(E)** Higher sensitivity in detecting short non-coding RNA in the small fraction of scComplete-seq.

Certain histone gene expressions were reported to vary according to cell cycle^8^. However, the expressions of these non-polyadenylated transcripts were undetectable in conventional scmRNA-seq. To investigate the dynamics of histone genes over the cell cycle, we estimated the cell cycle phase of single cells in scComplete-seq by the expression of cell-cycle-specific mRNA. We observed that some histone genes like *H2AC15, H4C9*, and *H3C1* were upregulated during the S and G1 phases, but their expressions decayed during the G to M transition (Fig. 2b-c). As histones play a crucial role in the packaging of DNA, their timely biosynthesis during the growth and DNA synthesis phase is essential for the mitosis process. On the other hand, expressions of other histone genes were either invariant (e.g. *H2AZ1, H2AX*) or exhibited an opposite relationship (e.g. *CENPA, H1*.*0*) over the cell cycle (Fig. 2b-c). Interestingly, these set of histones were also detected in conventional scmRNA-seq, indicating that they were naturally polyadenylated^9^ (Supp. Fig. 2). Thus scComplete-seq revealed the intricate transcriptional processing of different histone genes that were important for the regulation of the cell cycle^10^.

Next, we investigated if scComplete-seq could provide additional information, particularly regarding cell-type specific non-coding RNAs compared to conventional scmRNA-seq. We identified the top 50 most significant non-coding differential expressed genes (DEG) from scmRNAseq and found that the vast majority of them (96%) coincided with the top 200 non-coding DEG in scComplete-seq. However, only 78% of the top 50 non-coding DEG in scComplete-seq were found in the top 200 non-coding DEG in the scmRNA-seq. The non-coding DEGs that were only found in scComplete-seq consisted of short non-coding RNAs such as snRNA, tRNA, and miscellaneous RNA (Supplementary Data 1).

We expected the sensitivity of detecting non-polyadenylated, short non-coding RNA could be even higher if we inspected the sequencing library prepared from the small fraction. As we compared the normalized gene expression of short non-coding RNA from the sequencing libraries derived from the long and short fractions, we detected higher diversity (Fig. 2d) and expression (Fig. 2e) in the short fraction (155 and 245 small non-coding RNA expressed at least 0.01 copies were detected in the long and short fraction respectively). Thus, we proceeded to further investigate the single-cell short non-coding transcriptome through the short fraction library.

As we examined the top 50 short non-coding DEG, the small fraction detected twice the number (20) of short non-coding RNA compared to the large fraction (10), as expected based on their enhanced sensitivity. 7 of the short non-coding RNAs identified in the large fraction were also identified in the top 50 short non-coding DEG of the small fraction (Supp. Fig. 3, Supp Data 1).

The differentially expressed short non-coding RNAs among the three cell types were mainly found in the tRNA (nuclear and mitochondria), snRNA, and miscellaneous RNA biotypes (Fig. 3a). tRNAs are the most variable biotype among the cells, with specific cell types overexpressing particular tRNA species. The tRNA for *Cys-GCA* and *Ala-AGC* were upregulated in HEK293T cells. Meanwhile, K562 upregulated a large set of tRNA expressions including many mitochondria tRNA, *Ala-TGC, Ala-CGC, Ala-AGC, Gly-GCC, Gly-CTG, Gly-TCC, Gly-CCC, Arg-TCT, Cys-GCA, Glu-TTC, Ile-AAT, Ser-AGA, Thr-TGT* and *Val-TAC*. MCF7 upregulates *Gly-TCC, Gly-CCC, Thr-TGT, Glu-TTC*, and *Val-TAC*. Due to the important role of tRNA in protein synthesis, their differential abundances in different cell types have been reported to regulate the translation of cell-type-specific proteins^11^. Interestingly, we observed simultaneous overexpression of multiple tRNAs corresponding to the same amino acid (e.g. Alanine and Glycine in K562). Alanine and Glycine are the two smallest amino acids with non-polar, aliphatic side groups. Some previous works have reported variable mitochondria to nuclear tRNA ratios and coordinated expression of tRNA decoding amino acids with similar chemical properties in distinct tissue types and proposed that this could lead to different amino acid availability in these cells^11^.

**Figure 3.**
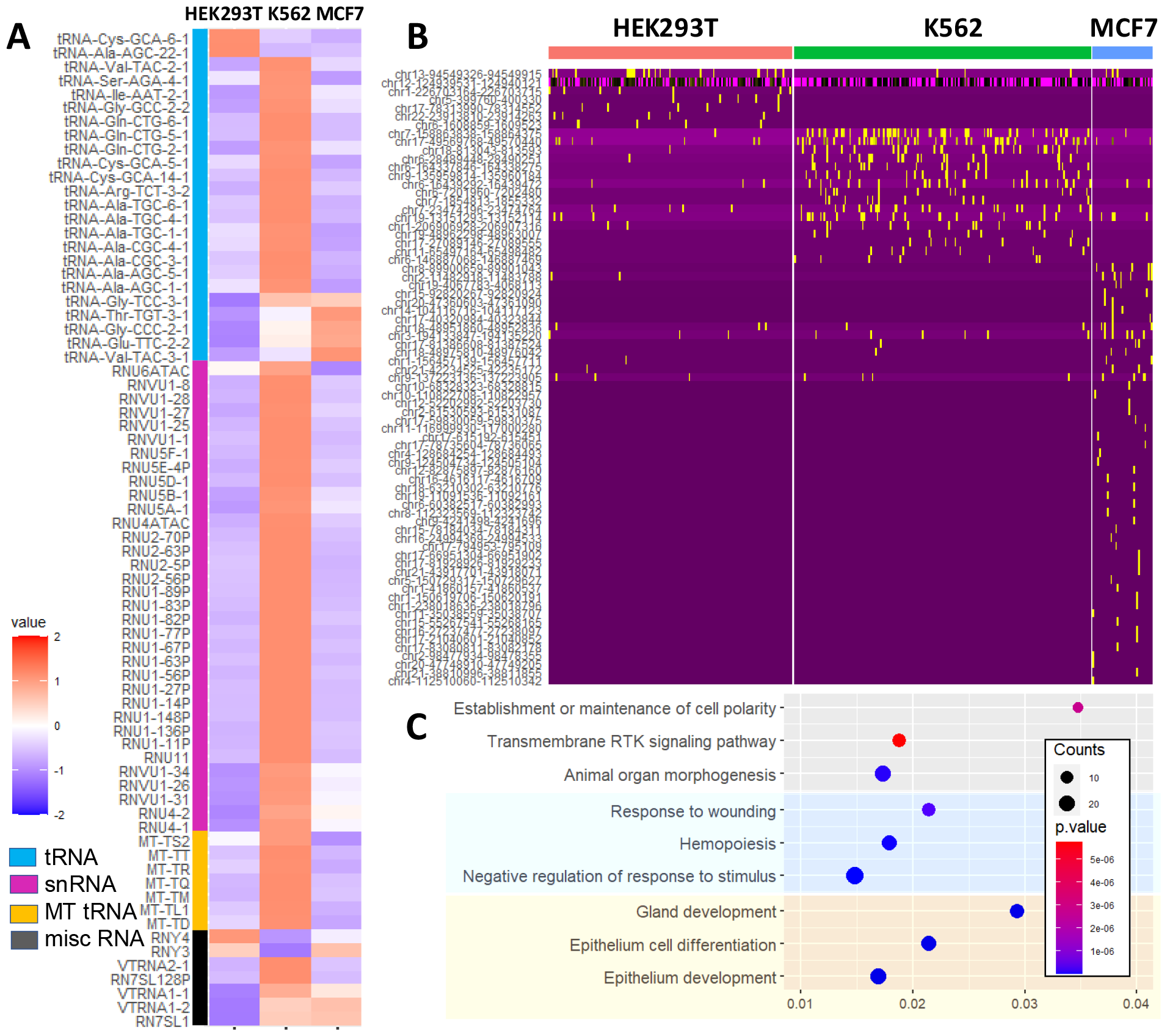
Short non-coding RNA and enhancer expression in the small fraction of scComplete-seq. **(A)** Differentially expressed short non-coding RNA in the HEK293T, K562, and MCF7 cell lines. **(B)** Enhancer expression in the HEK293T, K562, and MCF7 cell lines. **(C)** The top three GO terms (Biological Process) and the corresponding gene ratio for HEK293T (white background), K562 (blue background), and MCF7 (yellow background) cell lines. Gene counts and p values were shown by the size and color of the bubble respectively.

snRNAs are critical components of the spliceosomes that mediate the splicing of pre-mRNA. It was previously found that snRNA displayed distinct expression patterns across human cell types. With scComplete-seq the most pronounced observation was that K562 expressed the highest level of snRNA. This could be related to cell-type-dependent variations in global snRNA abundance, as observed in previous reports^12^. Meanwhile, HEK293T was observed to upregulate *RNU6ATAC* while MCF7 upregulates several snRNAs in the *RNVU1* and *RUU4* family. The differential expression of these snRNA suggested their possible roles in cell-type-specific splicing regulation. Lastly, in the class of miscellaneous RNA, we observed the upregulation of Y-RNA *RNY3* and *RNY4* in HEK293T and MCF7 cells. Vault RNA *VTRNA1* was upregulated in K562 and MCF7, but its counterpart *VTRNA2* was only overexpressed in K562. *RN7SL1*, the RNA component of signal recognition particle responsible for the insertion of secretory proteins into the lumen of the endoplasmic reticulum, was upregulated in K562 and MCF7.

Previous genome-wide studies have identified enhancers as common regions of extragenic transcription. As the expression of enhancer RNA (eRNA) corresponded to the activity of enhancers in the target gene, eRNA profiling would provide important information about gene regulation through enhancer interaction. Nonetheless, eRNAs are typically not detected in conventional mRNA sequencing approaches due to their low abundance and lack of polyadenylation. To investigate the potential for scComplete-seq to detect eRNA transcription, we counted RNA that mapped to enhancer sites derived from CAGE-seq^13^.

Compared to mRNA and other small non-coding RNA expression, eRNA expression was lower, consistent with its transient expression. Nevertheless, we could detect eRNA that were specific to HEK293T, K562, and MCF7 cells, suggesting different enhancer activities in these cell types. Comparison between the eRNA detected between HEK293T, K562, and MCF7 cells showed cell-type specific eRNA expression (Fig. 3b). Interestingly, HEK293T cells showed a reduced number of cell-type specific eRNA compared to K562 and MCF7 cells. To investigate the biological relevance of these cell-type-specific eRNAs, we obtained the putative target genes (nearest gene) of these enhancers and performed gene ontology (GO) analysis. The top two GO terms (Biological Process) for cell type-specific enhancers were *Response to Wounding* and *Hemopoiesis* for K562, *Epithelial Cell Differentiation* and *Gland Development* for MCF7, and *Establishment of Cell Polarity* and *Transmembrane Receptor Protein Tyrosine Kinase Signaling Pathway* for HEK293T respectively (Fig. 3c, Supplementary Data 2). These functional enrichments were consistent with the developmental origins of K562 as erythroleukemic cells, MCF7 as epithelial cells that originated from the mammary gland, and HEK293T as embryonic kidney cells. Thus, taken together, scComplete-seq provides comprehensive information on not only the mRNA variations but also the underlying transcriptional regulation mechanisms, in a high-throughput single-cell platform.

### scComplete-seq reveals cell-type specificity of coding and non-coding transcripts in PBMC

Our results thus far revealed the capability of scComplete-seq to perform comprehensive transcriptome profiling at single cell level. Specifically, the large fraction provides information regarding mRNA, lncRNA, and non-polyadenylated genes such as histones while the small fraction provides abundant information about the short non-coding RNA and enhancer RNA. As further proof of the concept of scComplete-seq to derive biological insights from heterogeneous cell populations, we applied this methodology to profile peripheral blood mononuclear cells (PBMCs). PBMCs have been extensively studied with scmRNA-seq, but the cell-type specific non-coding RNA and their changes in response to external stimulation have not been studied due to the lack of high throughput methods.

We performed scComplete-seq on unstimulated PBMCs, as well as PBMCs stimulated by lipopolysaccharide (LPS) and phorbol 12-myristate 13-acetate (PMA)-Ionomycin (PI) to study the cell-type- and stimulant-dependent comprehensive RNA changes at the single cell level. Besides labeling cells with hashtag antibodies, the PBMCs were also stained with a panel of feature-barcode antibodies for cell surface phenotyping. Following the optimized method described above, we obtained the large fraction and small fraction of the PAP-treated single-cell libraries of all three conditions. We performed data integration of the large fraction library across different stimulation conditions using the Seurat package^14^ to facilitate comparison among different experiments. Upon data integration, we obtained six major clusters that were populated in all three conditions (Fig. 4a, Supp. Fig. 4). Focusing on the unstimulated PBMCs, we annotated the six major populations based on their respective marker genes, consisting of CD14 monocytes (*CD14, LYZ*), CD16 monocytes (*MS4A7, FCGR3A*), CD4 T cells (*LEF1, TCF7*), CD8 T cells (*GZMK, CXCR3*), NK cells (*GZMB, NKG7*) and B cells (*MS4A1, CD79A*) (Fig. 4b). We also separately validated the identities of these clusters by the antibody barcodes (Supp. Fig. 5). As expected, CD4 T cells expressed CD3, CD28, and CD4 markers while CD8 T cells expressed CD3, CD28, and CD8 markers. CD16 was found in NK cells and some monocytes. CD27, a marker for memory B cells and naïve CD4 T cells was observed in a subset of the CD4 T and B cell cluster. The cell surface protein expression was thus consistent with the cell assignment obtained.

**Figure 4.**
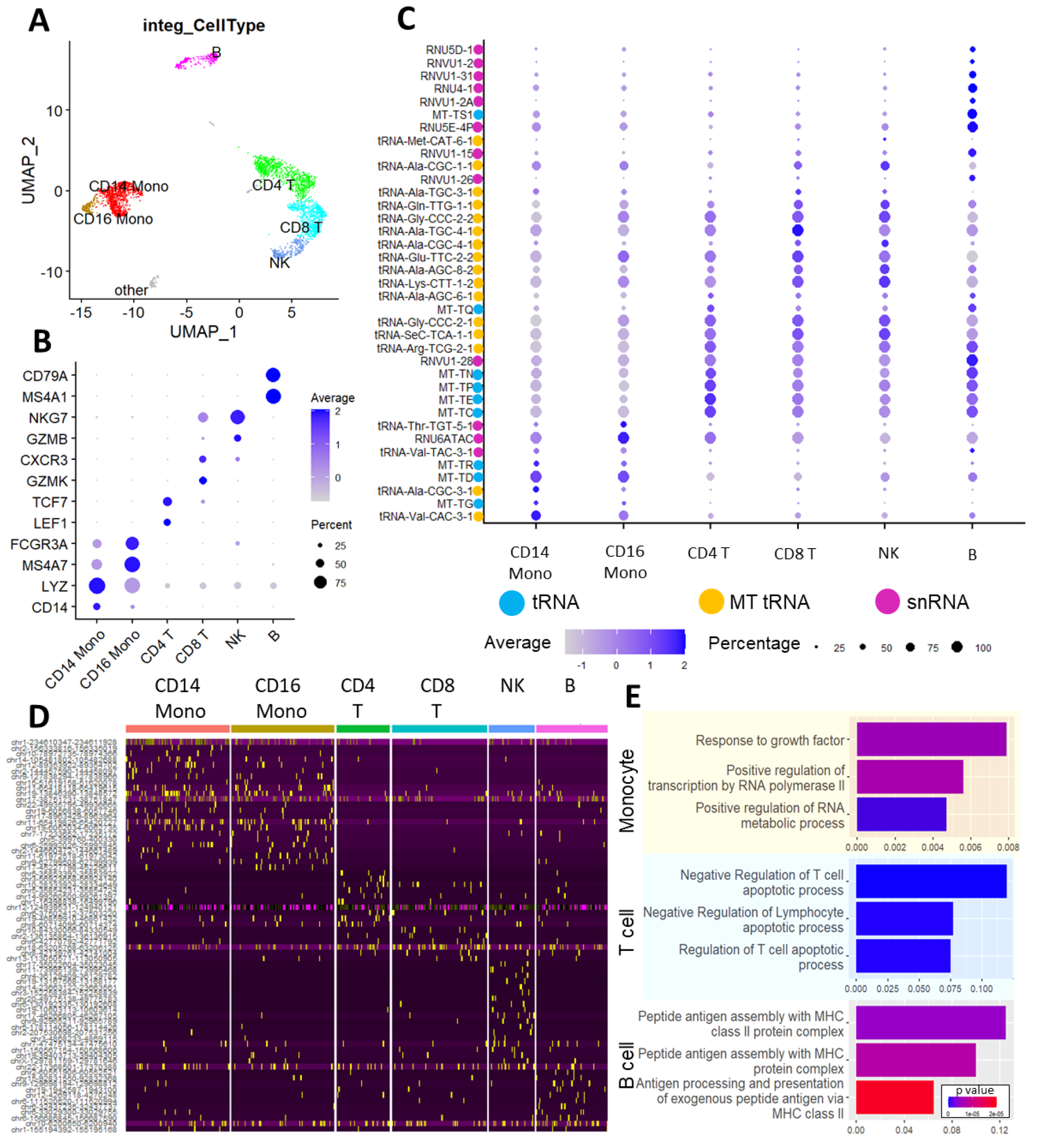
Cell-type-specific total transcriptome variations in PBMCs. **(A)** Different cell-type clusters were obtained in PBMCs with scComplete-seq. **(B)** Marker genes that defined the identities of cell clusters in PBMCs. **(C)** Differential short non-coding RNA expression in subpopulations of PBMCs. **(D)** Differential eRNA expression in the PBMC cell clusters. **(E)** The top three GO terms (Biological Process) and the gene ratio of the eRNA target genes for monocyte (yellow background), T cell (blue background), and B cell (white background). P values are shown by the color of the bar.

The mRNAs that were significantly overexpressed in each of the major clusters under non-stimulated conditions were examined (Supp. Fig. 6). Known T cells associated genes like LTB, TCF7, IL7R, CCR7, SELL, GZMK, CCl5, and CST7 were found to be highly expressed in the CD4 and CD8 T cells^15^. Both CD8 T cells and NK cells are cytotoxic effectors of the immune system, and they were found to share similar upregulation in genes like CST7^16^, NKG7, CXCR3, EOMES, APOBEC3G, PRF1, and KLRD1. However, NK cells also expressed some unique genes such as CEBPD, GNLY^17^, GFGBP2, CTSW^18^, and GZMB^19^. CEBPD was shown to promote NK-cell development and immunity^20^. In monocytes, high expression of LYZ, CTSB, CYP1B1^21^, MAFB^22^, S100A8, THBS1 were found in CD14 monocytes, while high expression of FGL2, MS4A7, CD74, CXCL16, HLA-DPA1, SERPINA1, TNFSF10 and VMO1 were found in CD16 monocytes respectively. B-lineage-associated genes like CD79A and CD79B were expressed in the B cells cluster. Some of the B cells expressed LTB and SELL, markers associated with mature B cells^23^.

Next, we examined the differential short non-coding RNA expression corresponding to the major cell types in PBMC (CD4 T, CD8 T, CD14 monocytes, CD16 monocytes, NK, and B cells. We obtained the bubble plot (Fig. 4c) and heatmap (Supp. Fig. 7) of the short non-coding RNA expression corresponding to the major cell types. There were a few interesting findings. Firstly, the small ncRNA profiles reflected the lineage relationship among the major cell types monocytes have a myeloid origin, while T cells, B cells, and NK cells originated from lymphoid progenitors. Accordingly, the small ncRNA profiles were most distinct between monocytes and other lymphoid cell types. Notably, a set of tRNA (*Thr-TGT-5-1, Ala-CGC-3-1, Val-CAC-3-1*) and mitochondria tRNA (*MT-TD, MT-TL2, MT-TG, MT-TR*) were upregulated in monocytes. The small nuclear RNA *RNU6ATAC* was upregulated in CD16 monocytes while the *7SK* small nuclear RNA was abundant in CD14 monocytes. Meanwhile, the cytotoxic immune cells, namely the CD8 T and NK cells, shared very similar small non-coding RNA profiles while the helper CD4 T cells were characterized by an upregulation of many mitochondria tRNA (*MT-TA, MT-TN, MT-TQ, MT-TS2, MT-TE, MT-TC, MT-TP*). Finally, B cells exhibited significant overexpression of numerous small nuclear RNA and their variants (*RNU*s and *RNVU*s) compared to other cells in PBMCs. Given that small nuclear RNAs are essential components in the spliceosome, our results could suggest the enhanced regulation of splicing activities in B cells such as in generating different antibody isotypes (e.g. membrane-bound IgM vs secreted IgG)^24^.

Following our previous demonstration of eRNA profiling, we next asked if the different cell types within PBMCs have distinct enhancer activities. Indeed, the eRNA profiles showed interesting patterns of eRNA patterns. The monocytes (CD14 and CD16) showed distinct eRNA profiles compared to the lymphocytes (CD4, CD8, NK, B). Interestingly, while the CD8 T cells were more similar to NK cells in small non-coding RNA, CD8 and CD4 cells were closer in eRNA expression. CD8 T cells were similar to NK cells in their cellular functions (cell-mediated cytotoxicity), but they were closer to CD4 T cells in terms of cell lineage. Hence, the different aspects of non-coding RNA profiled by scComplete-seq could reflect the distinct developmental origins (eRNA) versus functions (small ncRNA) of the cells. We further investigated the biological significance of the cell-type-specific enhancers and found that the top two GO Biological Process terms were *Negative Regulation of T Cell Apoptotic Process* and *Negative Regulation of Lymphocyte Apoptotic Process* in T cells, *Peptide Antigen Assembly with MHC Class II Protein Complex* and *MHC Class II Protein Complex* in B cells, and *Response to Growth Factor* and *Positive Regulation of Transcription by RNA Polymerase II* in monocytes (Supplementary Data 2). Our results revealed important biological insights into enhancer-regulated gene networks that were important for the maintenance of T cells, class II-restricted antigen presentation in B cells, as well as transcriptional activation in monocytes.

### Deciphering total transcriptome changes in PBMCs under stimulation

Our results showed that PBMCs were not only diverse in protein-coding genes, but the different cell types were also very distinct in noncoding RNA. An interesting property of PBMC is their ability to respond distinctively to different stimuli. For example, the combination of PMA and ionomycin (PI) mimics antigen engagement in the adaptive immune cells and stimulates their effector functions, while exposure to LPS stimulates the innate immune response through TLR signaling^25^. While clear mRNA expression changes have been reported for this stimulation, we expect that concomitant changes in the non-coding RNA would shed light on the molecular responses of different cell types. To investigate this, we examined the comprehensive transcriptome changes through scComplete-seq for PBMCs stimulated by PMA-Ionomycin or LPS.

We first performed differential gene analysis for mRNA for all cell types between stimulated (PI or LPS) and unstimulated conditions. PMA activates protein kinase C, while ionomycin is a calcium ionophore and stimulation with these compounds bypasses the T cell membrane receptor complex and will lead to the activation of several intracellular signaling pathways, resulting in T cell activation and production of a variety of cytokines^26^. As expected, we observed upregulation of Th1 cytokines such as *IL2, TNF* and *IFNG* in the PI-treated T cells (Fig. 5a). On the other hand, LPS, the principal component of the outer membrane of Gram-negative bacteria, is a potent stimulator of monocytes, resulting in production of an orthogonal set of cytokines (*IL1B, IL6, IL10*) that are essential for innate immune response (Fig. 5b).

**Figure 5.**
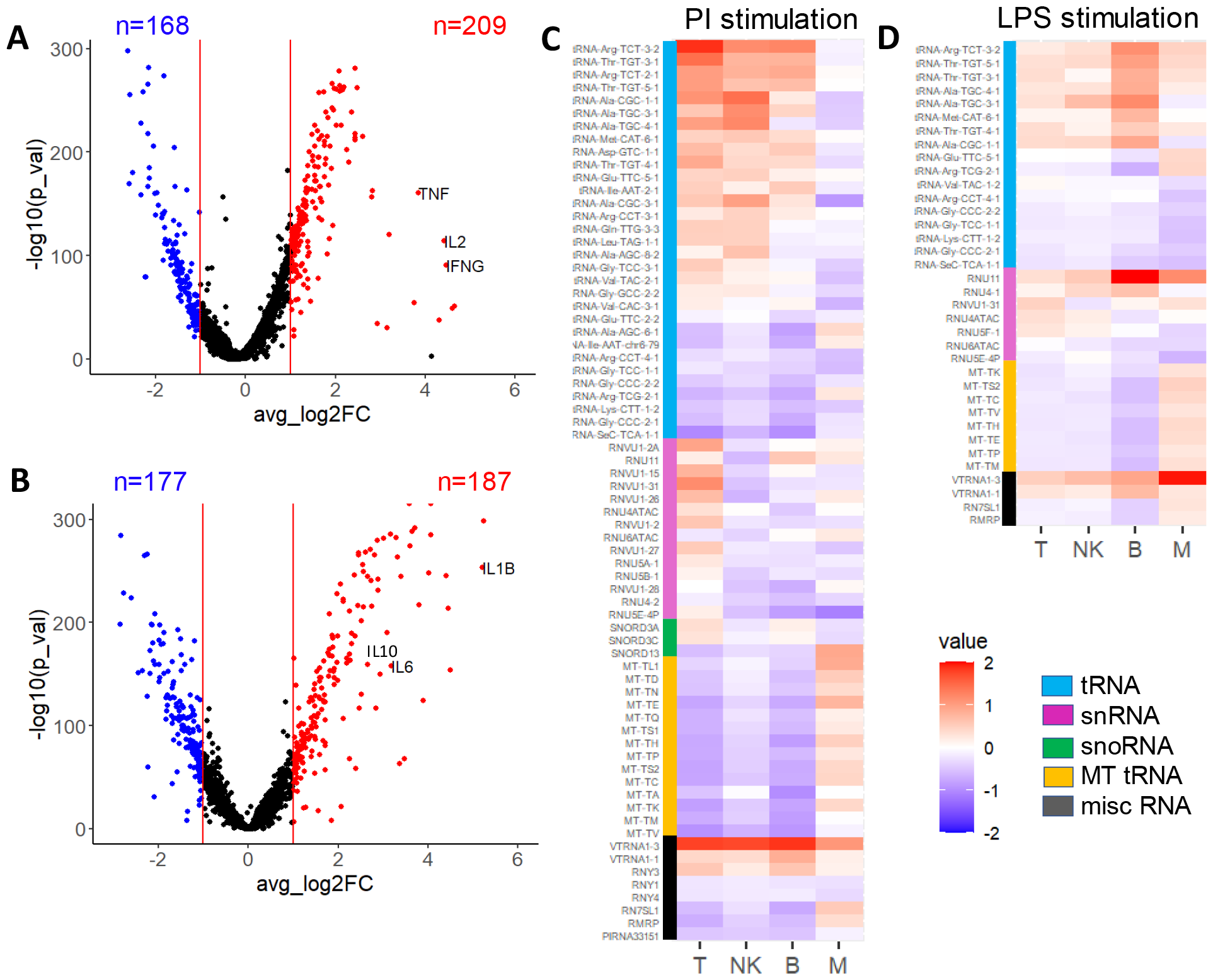
Differential coding and non-coding gene expression upon PI or LPS stimulation. Volcano plot showing gene expression changes in (A) T cells upon PI stimulation and **(B)** monocytes upon LPS stimulation. Heatmap of the differentially expressed short non-coding RNA in different cell types upon **(C)** PI and **(D)** LPS stimulations compared to unstimulated subsets.

Next, we proceeded to identify changes in small non-coding RNA in each cell type upon stimulation. The T cell subpopulations (CD8 and CD4) and monocyte subpopulations (CD14 and CD16) were combined in the analysis to focus on the changes in the major cell types. During PI stimulation, there were significant changes in the expression of 69 small non-coding RNA among the different cell types (Fig 5c). Arranging them by their biotypes (tRNA, MT-tRNA, snRNA, snoRNA, misc RNA), there were several interesting observations. First, monocytes responded to PI stimulation in a very different manner compared to the lymphocytes (T, B, NK). Monocytes respond by upregulating many mitochondria tRNA while lymphocytes respond differently by upregulating cytoplasmic tRNA. Previous studies have shown that compared to lymphocytes, monocytes are more sensitive to oxidative stress. Oxidative stress induced mitophagy in monocytes during inflammation which needs to be compensated by increased mitochondria biogenesis^27^. Enhanced mitochondria biogenesis may explain the upregulation of mitochondria tRNA in monocytes upon stimulation. Within the lymphocytes, T cells demonstrated the most apparent small non-coding RNA changes upon PI stimulation, as expected since PMA selectively activates PKC, a core protein in the T cell immunological synapse. Notably, a set of snRNA typically associated with the spliceosome complex was upregulated in T cells compared to other lymphocytes. Alternative splicing of RNA to generate protein isoforms with different functions during T cell activation has been reported – the most well-known example being alternative splicing of the transmembrane phosphatase CD45 to yield the shorter isoform CD45RO with reduced phosphatase activity^28^. Our results shed light on the set of non-coding RNAs that may play important roles in this post-transcriptional regulation.

Meanwhile, examination of small ncRNA changes during LPS stimulation yielded similar general trends (Fig 5d). Upregulation of many MT-tRNA was similarly observed in monocytes, but we also observed upregulation of many cytoplasmic tRNAs in this case. Accordingly, monocytes experienced the most apparent small ncRNA among all cell types in LPS stimulation. The three populations of lymphocytes shared similar trends of small ncRNA changes. However, the most significant change within the lymphocyte populations was seen in B cells instead of T cells. B cells possess TLR4, the receptor for LPS. LPS can stimulate dual receptor signaling by bridging the B cell receptor and Toll-like receptor 4 (BCR/TLR4)^29^, thus leading to relatively large changes in small ncRNA in B cells.

We also examined enhancer RNA changes in the cell populations upon stimulation. Distinct eRNA expression changes were observed in all cell types, but the degree of changes varied. The greatest eRNA changes were found in monocytes for LPS stimulation and T cells for PI stimulation. Thus, we next focused on investigating the biological implications corresponding to these two cell types for both stimulations. The top two GO term enrichment were *Response to Cytokine* and *Regulation of Cell Adhesion* in monocytes stimulated with LPS, *Response to Oxidative Stress* and *Cell Activation* in monocytes stimulated with PI, and *Mononuclear Cell Differentiation* and *Regulation of Lymphocyte Activation* in T cells stimulated with PI (Fig. 6c, Supplementary Data 3). No GO term enrichment was found for T cells stimulated by LPS due to the insignificant changes in eRNA. Our results highlighted that the same cell type (e.g. monocyte) engaged with different biological processes upon different stimulations (e.g. oxidative stress in PI and cytokine response in LPS). Secondly, under the same stimulation (e.g. PI), the different cell types responded differently (e.g. oxidative stress for monocytes and mononuclear cell differentiation for T cells).

**Figure 6.**
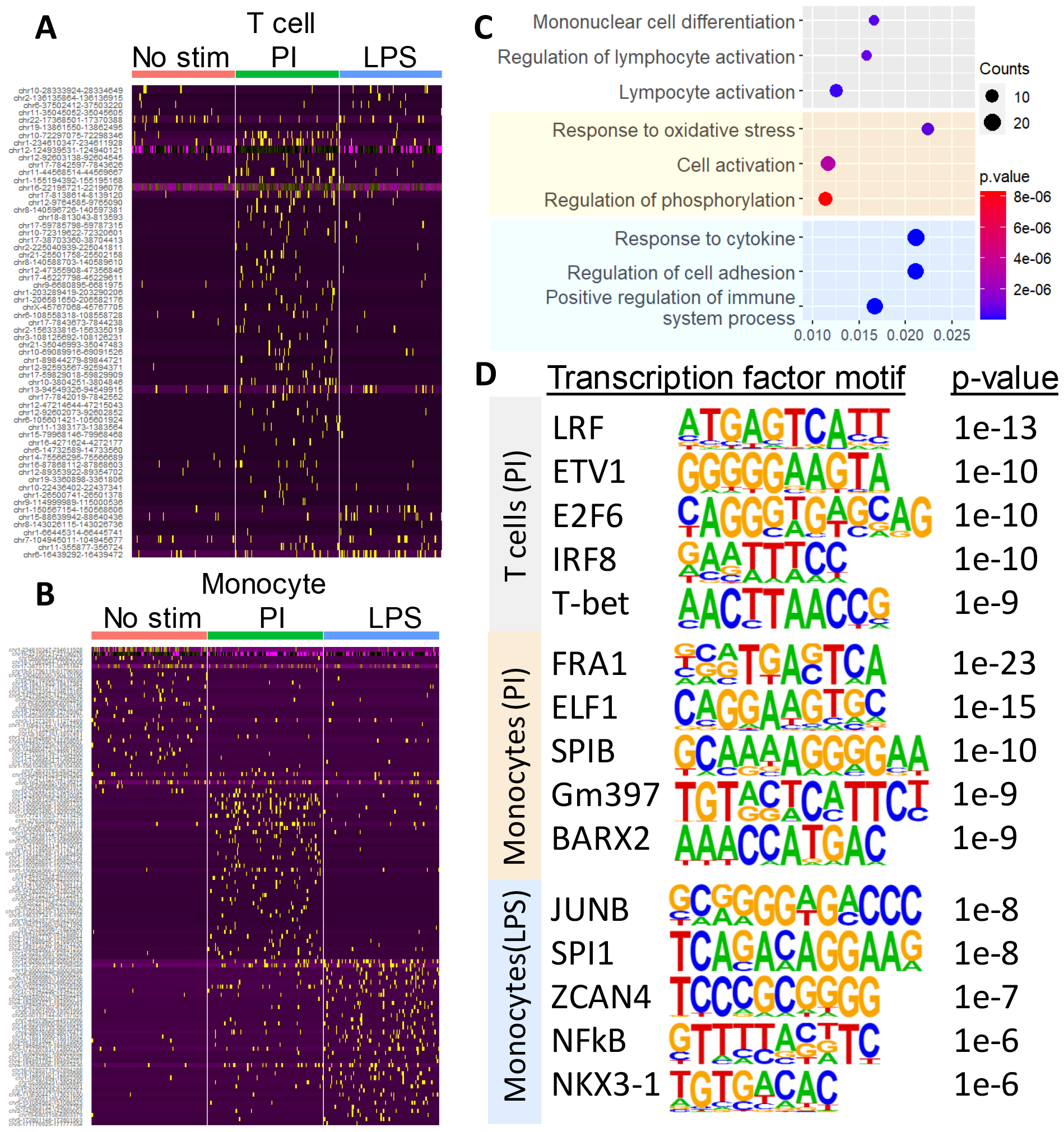
Changes in eRNA profiles during PI and LPS stimulation. Differential enhancer expression during PI and LPS stimulation in **(A)** T cells and **(B)** monocytes. **(C)** The top three GO terms (Biological Process) and the corresponding gene ratio for T cells stimulated with PI (white background), monocytes stimulated with PI (yellow background), and monocytes stimulated with LPS (blue background). Gene counts and p values are shown by the size and color of the bubble respectively. **(D)** Transcription factor motifs in the differential eRNA and their corresponding p-values obtained from T cells stimulated with PI (white), monocytes stimulated with PI (yellow), and monocytes stimulated with LPS (blue) respectively.

Finally, the activation of different enhancers is often associated with the binding of various transcription factors. Given that scComplete-seq allowed us to identify enhancer usage at single-cell resolution, we could potentially identify transcription factor motifs on the stimulation-induced enhancers (Fig. 6d). Applying a motif-finding program on the identified stimulation-induced eRNAs, we found that JUNB, a transcription factor regulating gene activity following growth factor response^30^ was the top motif for monocytes stimulated by LPS. NFKB, a central regulator of inflammation^31^ was also identified within the top 5 motifs. FRA1, a protein that can dimerize with proteins of the JUN family forming the transcription factor complex AP-1 and is essential for monocyte-macrophage differentiation^32^, was identified as the top motif for monocytes stimulated by PI. Another noteworthy motif was ELF1, a protein primarily expressed in lymphoid cells that can act as both an enhancer and a repressor to regulate the transcription of various genes^33,34^. Finally, LRF was the top motif for T cells stimulated by PI. LRF has been reported to be an essential transcription factor for T helper cell differentiation^35^. TBET, the master regulator of type 1 immune response in T cells^36^ was also identified. Thus, the ability of scComplete-seq to identify eRNA at the single-cell level allowed the prediction of direct transcription factor binding at enhancers, which was previously impossible.

## DISCUSSION

Today, single-cell transcriptomic analyses have found applications in almost every field of cell biology, ranging from mapping the human cell atlas to uncovering subpopulations of cells associated with diseases. Nonetheless, conventional high-throughput scmRNA-seq platform technologies are focused on polyadenylated RNA and often miss important information that is found in the non-polyadenylated RNA species. Although there have been several recent efforts to enable the profiling of non-polyadenylated RNA in single cells, a truly automated high-throughput assay that is generally accessible to the scientific community is notably absent.

Here, we describe scComplete-seq, a high-throughput single-cell analysis platform capable of providing comprehensive (both polyadenylated and non-polyadenylated) transcriptome information that is compatible with a widely accessible automated single-cell processing instrument. By combining multiple reactions in a single step and substituting incompatible reagents with alternatives, the entire workflow only involves the use of a single commercially available droplet microfluidic chip instead of requiring multiple custom chip operations. This is an important breakthrough that would enable even non-technical specialists to access single-cell whole transcriptome sequencing easily.

A key innovation of scComplete-seq is the capability to preserve both the long (majority coding) and short (majority noncoding) RNA in single cells and link them to their common cell barcode. Processing the long RNAs with a cDNA fragmentation approach enables 3’ end counting for accurate gene expression with low sequencing depth. Meanwhile, short RNA recovered from the complementary small fraction does not undergo fragmentation which enriches whole transcripts and improves their mapping and alignment. Using this approach, we show that scComplete-seq not only retains the information available from conventional scmRNA-seq but also provides insights into the heterogeneity of non-polyadenylated long RNA (e.g. histones), short non-coding RNA (e.g. tRNA, snRNA, miscRNA) and enhancer RNA.

There are yet a few areas that can be further improved on scComplete-seq. Firstly, while other methodologies such as SMART-seq Total, which employs polyadenylation and template switching, have successfully recovered very short small non-coding RNA such as miRNA and piRNA, we observed notable depletion of them in scComplete-seq. We attribute this to their inadvertent loss due to overzealous adapter removal in our library preparation process. The use of more fine-tuned procedures such as BluePippin may remove the majority of adapters with less depletion of very short RNA species. Secondly, lowly expressed transcripts such as eRNA are not adequately profiled in cells with low sequencing depths, and we have to filter off cells with lower read counts to get a better picture of the eRNA profile. Higher-depth sequencing of the small fraction scComplete-seq library would be recommended to investigate eRNA heterogeneity in greater detail. Finally, even though we have effectively removed the most abundant rRNA species, there are a few highly abundant species such as Vault RNA that make up a significant proportion of the sequenced reads. Additional CRISPR-mediated removal^37^ of these abundant sequences may allow us to further improve the effectiveness of scComplete-seq.

In a proof of concept, the application of scComplete-seq on resting and stimulated PBMCs revealed diverse patterns of snRNA and tRNA expression dependent on cell type and stimulation. Given the roles of snRNA on splicing and tRNA on protein translation, scComplete-seq may provide a further indication of the degree of post-transcriptional processing that would shape the cell phenotype. Additionally, we showed that scComplete-seq could detect eRNA expression, which generally correlates with enhancer activation and can serve as a direct marker of active enhancers^38,39^. GO analysis of the putative genes targeted by these enhancers showed enrichment in biological processes consistent with these cell types while further analysis of these eRNA revealed distinct transcription factor motifs that could explain the mechanisms of gene regulation. Thus, scComplete-seq may provide a critical link to epigenetic regulation (through small RNA and enhancers) of gene expression in single cells. We envision that scComplete-seq will be a convenient but powerful toolkit to explore the comprehensive transcriptome at the single-cell level.

## SUPPLEMENTARY INFORMATION

Supplementary information is available for this manuscript.

## ACKNOWLEDGEMENTS

This work was supported by funding from the Singapore Ministry of Education Academic Research Fund Tier 2 (MOE-000063) and the Institute for Health Innovation and Technology (iHealthtech), National University of Singapore (NUS).

## AUTHOR CONTRIBUTIONS

F.B.D and L.F.C conceived and designed the study. F.B.D. and S.W.Y.N. performed the experiments. F.B.D., R.Z.T., and L.F.C. performed data analysis. F.B.D., R.Z.T., and L.F.C. wrote the manuscript. All authors approved the submission.

## MATERIALS AND METHODS

### Cells and reagents

This study was approved by the National University of Singapore Institutional Review Board (NUS-IRB no. H-18-038E). Residual apheresis blood samples were obtained from anonymous healthy donors who have given consent for their blood to be used for research at the Health Science Authority of Singapore. Ficoll-Paque Plus (GE Healthcare, 17-1440-02) was used for PBMC separation. RPMI 1640 medium (Life Technologies, A1049101) with 10% fetal bovine serum (Life Technologies, 1027016 and 1% Penicillin-Streptomycin (Life Technologies, 15070063) was used in cell culture. Human breast cancer cell line MCF7, human chronic myelogenous leukemia cancer cell line K562 and human embryonic epithelial kidney cell line HEK293T were cultivated in Dulbecco’s Modified Eagle Medium with high glucose (#11965084, Life Technologies) supplemented with 10% fetal bovine serum Life Technologies, 1027016) and 1% Penicillin-Streptomycin (#15070063, Life Technologies) at 37 °C and 5% CO2. Adherent cell lines (MCF7 and HEK293T) were passaged at 80% confluence every 3–4 days, whereas K562 was split 1:7 every 5 days.

TotalSeq-A antibodies used for staining PBMCs are listed in Supplementary Table 1. DNA oligos were synthesized by Integrated DNA Technologies and their sequences are listed in Supplementary Table 2. For immunostaining, Fc-blocking reagent (BioLegend, 422301) was used to block Fc receptor-introduced nonspecific staining. Phosphate-buffered saline (PBS) with 0.02% FBS was used as the staining and washing buffer.

### Cell preparation for scComplete-seq

For scComplete-seq of cell lines, MCF7 and HEK293T cells were harvested with 1X TrypLE (Gibco, 12563011) treatment, pelleted at 350g for 3 min and the supernatant was aspirated. After aspiration of the supernatant, the cells were washed in a washing buffer. Phosphate-buffered saline (PBS) with 0.02% FBS was used as the staining and washing buffer. K562, MCF7, and HEK293T cells were then blocked with Fc-blocking agent (BioLegend, 422301) at 4 °C for 30 min and labeled with sample identifier hashtags (0.5 µg each TotalSeq-A anti-human Hashtag per million cells. Cells were washed with wash buffer 3 times and pooled together (equal number of cells each, loading 4800 cells in total.)

For scComplete-seq of PBMCs, ∼10 million PBMCs were thawed at 37 ºC immediately after removal from liquid nitrogen. Cells were rested at 37 ºC for 2 h in RPMI Complete Media (RPMI 1640 medium (Life Technologies, A1049101) with 10% fetal bovine serum (Life Technologies, 1027016) and 1% Penicillin-Streptomycin (Life Technologies, 15070063) with 5% CO2 at 37 °C incubator. The cells were seeded at ∼2 million per well in 6 well plates in three conditions: 1) Unstimulated, 2) 2 µl/mL PMA/Ionomycin (1x final dilution, eBioscience Cell Stimulation Cocktail 500X, 00-4970-93), and 3) 0.02 mg/mL LPS (4x final dilution, eBioscience Lipopolysaccharide Solution 500X, 00-4976-93) for 8 h in RPMI Complete Media. The cells from different conditions were then blocked with Fc-blocking agent and labeled with sample identifier hashtags (0.5 µg each TotalSeq-A anti-human Hashtag per million cells) and Fc-blocking agents at 4 °C for 30 min. Cells were washed with wash buffer 2 times and pooled together (equal number of cells per condition, and loading 16,000 cells in total). An Ab-oligo cocktail (0.125 µg each TotalSeq-A antibody per million cells) was prepared in 45 µL of staining buffer and 5 µL of Fc-blocking reagents immediately before use. The pooled barcoded cell samples were stained with surface marker Ab-oligo cocktail at 4 °C for 40 min. Cells were washed with wash buffer 3 times and resuspended in PBS.

### Implementation of scComplete-seq in automated high-throughput single-cell platform

scComplete-seq was performed with a slight modification of the 10x Genomics single cell 3′ reagent kits (v3.1 Chemistry) and protocol. The RNA-TSO was replaced with LNA-TSO. PAP enzyme and ATP (NEB, M0276L) were also added to the cell mix to facilitate *in vitro* polyadenylation. The final modified reagent mix (75 µL) consists of 18.8 µL RT Reagent B, 2 µL Reducing Agent B, 8.7 µL RT Enzyme C, 3 µL of LNA-TSO (100 µM), 3 µL of PAP enzyme (50 U/µL), 18 µL of ATP (10 mM) and 21.5 µL of cells.

After loading the cells and reagents into the single-cell chip (Chromium Next GEM Chip G Single Cell Kit) and instrument, the single-cell encapsulation process was performed. After cell encapsulation in droplets, they were collected and incubated according to a modified protocol of 25 min at 37 ºC, 45 min at 53 ºC, and 5 min at 85 ºC. The additional incubation at 37 ºC allowed *in vitro* polyadenylation reaction to be carried out.

The droplets were demulsified followed by 10X Genomics protocol using magnetic beads. The recovered material includes complementary DNA (cDNA) of total RNA, surface marker barcodes, and sample hashtags. After sample clean-up using silane beads, 30 µL of the eluted DNA samples were mixed with 7 µL of TSO digestion mix (3.5 µL UDG-DNA glycosylase and 3.5 µL UDG Buffer, NEB M0280S) and incubated at 37 ºC for 30 min, followed by 85 ºC for 5 min to remove unreacted TSO. 37 µL of the sample was mixed with 50 µL of 2X Amp Mix (Chromium Next GEM Single Cell 3’ Library kit v3.1) to obtain a final 100 µL solution containing 1.5 µM of forward preamplification primer, 1.5 µM of reverse preamplification primer and 0.005 µM of HTO additive primer. Subsequently, the sample was pre-amplified using a thermal protocol of 98 ºCfor 3 min, 12 cycles of 98 ºC for 15 s, 63 ºC for 20 s, and 72 ºC for 1 min, and a final extension step at 72 ºC for 1 min.

The pre-amplified products were divided into 4 aliquots. In one aliquot, the PCR product was cleaned up and followed by the 10X Genomics double size selection step using SPRIselect reagent (Beckman Coulter) to separate the protein library (2X SPRI) from the large cDNA library (0.6X SPRI). The amplified cDNA product was subjected to the large fraction gene expression library construction according to the kit instructions. Part of the protein library product (5 µL) was used for the preparation of sequencing libraries corresponding to the sample hashtags, and an additional part (5 µL) was used for the preparation of sequencing libraries corresponding to surface markers (PBMC experiment only). In another aliquot of pre-amplification product, the PCR product was size selected using SPRIselect reagent to recover the small cDNA (1X SPRI).

Library preparations for the small fraction cDNA, ADT, and HTO were performed in 1X KAPA HiFi PCR reagent (Roche) with 1 µM of forward primer and 1 µM of reverse primer and thermal protocol as specified below. For scmRNA-seq and scComplete-seq long fragments (0.6X SPRI), the preamplification sample is fragmented, end-repaired, A-tailed, and indexed library was prepared following 10X Chromium v3.1 protocol. For the small fraction cDNA, we used the library forward primer and indexed reverse primer and the following thermal protocol: 95 ºC for 3 min; 11/12 cycles (PBMC/Cell Lines) of 98 ºC for 20 s, 65 ºC for 30 s and 72 ºC for 1.5 min; and a final extension step at 72 ºC for 3 min. For the ADT library, we used the library forward primer and indexed reverse primer and the following thermal protocol: 95 ºC for 3 min, 15 cycles of 98 ºC for 20 s, 60 ºC for 30 s, 72 ºC for 20 s, and 72 ºC for 5 min for the final extension. For the HTO library, we used the library forward primer and indexed reverse primer and the following thermal protocol: 95 ºC for 3 min, 15 cycles of 98 ºC for 20 s, 64 ºC for 30 s, 72 ºC for 20 s, and final extension of 72 ºC for 5 min.

cDNA corresponding to rRNA in both the large and small size cDNA fractions were removed using the SEQuoia kit (Biorad, 17006487) according to the manufacturer’s recommendation. The size and concentration of final library constructions were verified using Agilent 2100 Bioanalyzer.

### Data processing

The libraries for cDNA (large and small fractions), sample hashtag barcodes, and surface protein barcodes were indexed, pooled, and paired-end (2x 150bp) sequenced in the Illumina NovaSeq 6000 platform. As the cDNA in scComplete-seq is expected to include many other species beyond conventional mRNA-sequencing, we generated a custom GTF file by merging the Ensembl Human Reference with external small non-coding RNA annotated in DASHR v2 and miRNA using the MGCount approach^40^. We also appended the putative eRNA annotations obtained from CAGE-seq^13^ in the custom GTF file. The details of the annotation file are provided in Supplementary Table 3. The software and computational tools used in the analysis are given in Supplementary Table 4.

Standard pre-processing was employed for all single-cell RNA-seq datasets. Cutadapt^41^ (v4.4) with parameters -u 3 -m 16 -a AAAAAAAAAAAAAAA, and riboDetector^42^ (v0.2.7) with parameters -l 147 -e rrna --chunk_size 256 were used for rRNA removal and reads trimming as outlined in Supplementary Figure 8. Trimmed cDNA FASTQ data were processed for cell barcodes calling and unique molecular identifier (UMI) counting of the cDNA using STARSolo^43^ (v2.7.10b) with modified parameters (--soloType CB_UMI_Simple --soloStrand Forward -- outFilterMultimapNmax 50 --soloCBwhitelist celllranger3M-february-2018.txt --soloUMIlen 12 --outFilterScoreMinOverLread 0 --outFilterMatchNminOverLread 0 –winAnchorMultimapNmax 100 --outFilterMatchNmin 15 --outFilterScoreMin 14 --outFilterMismatchNoverLmax 0.05 -- seedMultimapNmax 100000 --seedSplitMin 9 --seedSearchStartLmax 25 --soloCellFilter EmptyDrops_CR --soloCBmatchWLtype 1MM_multi_Nbase_pseudocounts --soloUMIdedup 1MM_CR --soloFeatures Gene GeneFull SJ Velocyto --soloMultiMappers EM) using the custom GTF. Count data were retrieved for the following gene biotypes: protein coding, lncRNA, antisense, IG LV gene, IG V gene, IG V pseudogene, snRNA, IG D gene, IG J gene, IG J pseudogene, IG C gene, TR V pseudogene, TR D gene, TR J gene, TR J pseudogene, TR C gene, pre-miRNA, vault RNA, misc RNA, MT rRNA, MT tRNA, ribozyme, rRNA, sRNA, scaRNA, siRNA, piRNA, tRNA, miRNA, scRNA, snoRNA and eRNA. The generated cell barcodes were used to find individual cell barcode-matched expression profiles of sample hashtags and surface proteins using CITE-seq-Count (v1.4.0) with a Hamming distance of 1 for both cell barcode calling and UMI counting.

The raw count matrices were imported into the Seurat package (v4.0), and dead cells were removed by filtering out cells expressing >10% mitochondrial transcript counts for cell lines and >20% for PBMCs. For the analysis of cell lines, we chose cells that had RNA counts >500 and total features >200. For the analysis of PBMCs, we chose cells that had RNA counts between 1000-10000 and total features between 350-2500. The respective thresholds were determined based on the inflection points of the distribution of these features for sample type. Data normalization and scaling were implemented using the NormalizeData and ScaleData functions in Seurat. Centered-log transformation (CLR) was applied to sample hashtags and surface markers.

In the analysis of cell lines, the cell identity was determined from the cell hashtag with the highest signal^44^. For cell cycle phase assignment, each cell was assigned into discrete classes (G2M/G1/S) based on its expression of G2/M and S phase markers using the CellCycleScoring function in Seurat.

To analyze the PBMC, the large fraction of cDNA data was first normalized and the top 2000 variable features were determined. Anchors were then built using the large fraction library of the non-stimulated, PI-stimulated, and LPS-stimulated cells. 5 neighbors were used to pick anchors, 200 neighbors were used to filter anchors, and 30 neighbors were used to score anchors. Finally, the datasets were integrated using the pre-computed anchors using the top 20 dimensions. This analysis was implemented using the FindIntegrationAnchors and IntegrateData functions in Seurat. Next, we performed PCA dimensionality reduction on the integrated dataset, constructed a shared nearest neighbor graph for k = 20 using the top 20 PCA dimensions and performed cluster determination using the Louvain algorithm with a resolution of 0.5. The analysis was implemented using the FindVariableFeatures, RunPCA, FindNeighbors, and FindClusters functions in Seurat. Through these analyses, six major clusters were populated in all three conditions. The identities of these clusters were annotated based on the expression of known marker genes of cell types. All mRNA analyses were performed on the large fraction cDNA datasets while small non-coding RNA and eRNA analyses were performed on the small fraction cDNA datasets.

### Identification of differentially expressed genes

Differential gene expression between different cell populations was performed in Seurat using the Wilcoxon rank-sum test. To identify differential enhancer expression between different cell lines, a minimum fraction of 0.01 and a threshold log fold change of 0.01 were used. To identify differential enhancer expression between subpopulations in PBMC, a minimum fraction of 0.2 and a threshold log fold change of 0.2 were used. The genes were then ranked by their p-values to determine the top significant genes. To identify differential enhancer expression between PBMC subpopulations, a minimum fraction of 0.04 and a threshold log fold change of 0.1 were used on cells with transcript counts greater than 2500. To identify differential enhancer expression between stimulated (PI or LPS) and unstimulated conditions in PBMC, a minimum fraction of 0.04 and a threshold log fold change of 0.1 were used.

### Gene Set Enrichment Analysis (GSEA) and identification of transcription factor motifs

Differentially expressed enhancer RNA between different cell types or different stimulation conditions were identified as above. We obtained the coding genes that are nearest to these enhancer RNAs using the *closest* function in *bedtools*. The lists of these genes were used as input to the Molecular Signature Database (MSigDB)^45^ to find enrichment in Gene Ontology (GO) biology processes. The GO terms were ranked by the gene ratio and GO terms with less than 5 genes were excluded in the PBMC stimulation analysis. Transcription factor motif analysis was performed using the HOMER^46^ software suite to identify de novo motifs enriched in the differentially expressed enhancer RNA.

### Data Availability Statement

Raw sequencing and processed data generated in this project have been deposited to the Gene Expression Omnibus (GEO) with the accession code GSE261371.

