## Supplementary Information for "Automated high-throughput profiling of single-cell total transcriptome with scComplete-seq"

**Supplementary Table 1:** List of antibodies used in this study.

| **Antibody** | **Clone** | **Company** | **Catalog number** |
|---|---|---|---|
| TotalSeq-A0251 anti-β2M | LNH-94 | BioLegend | 394601 |
| TotalSeq-A0252 anti-β2M | LNH-94 | BioLegend | 394603 |
| TotalSeq-A0253 anti-β2M | LNH-94 | BioLegend | 394605 |
| TotalSeq-A0034 anti-CD3 | UCHT1 | BioLegend | 300475 |
| TotalSeq-A0045 anti-CD4 | SK3 | BioLegend | 344649 |
| TotalSeq-A0148 anti-CCR7 | G043H7 | BioLegend | 353247 |
| TotalSeq-A0063 anti-CD45RA | HI100 | BioLegend | 304157 |
| TotalSeq-A0386 anti-CD28 | CD28.2 | BioLegend | 302955 |
| TotalSeq-A0154 anti-CD27 | O323 | BioLegend | 302847 |
| TotalSeq-A0156 anti-CD95 | DX2 | BioLegend | 305649 |
| TotalSeq-A0391 anti-CD45 | HI30 | BioLegend | 304064 |
| TotalSeq-A0146 anti-CD69 | FN50 | BioLegend | 310947 |
| TotalSeq-A0046 anti-CD8 | SK1 | BioLegend | 344751 |
| TotalSeq-A0083 anti-CD16 | 3G8 | BioLegend | 302061 |
| TotalSeq-A0047 anti-CD56 | 5.1H11 | BioLegend | 362557 |

**Supplementary Table 2:** Primers used in the study.

| DNA Oligo | Sequence | Application |
|---|---|---|
| LNA-TSO | /5Biosg/UGA CUG GAG UUC CUU GGC ACC CGA GAA UUC CArG rG+G | TSO |
| Reverse preamplification primer | GTG ACT GGA GTT CCT TGG CA | Preamp |
| Forward preamplification primer | CTA CAC GAC GCT CTT CCG ATC T | Preamp |
| HTO additive primer | GTG ACT GGA GTT CAG ACG TGT GCT CTT CCG AT*C* T | Preamp |
| Library forward primer | *AAT GAT ACG GCG ACC ACC GAG ATC T*AC ACT CTT TCC CTA CAC GAC GC*T*C | Short cDNA library, ADT library, HTO library |
| Library indexed reverse primer | *CAA GCA GAA GAC GGC ATA CGA GAT* [8bp i7 index] GTG ACT GGA GTT CCT | Short cDNA library, ADT library |
| HTO indexed reverse primer | *CAA GCA GAA GAC GGC ATA CGA GAT* [8bp i7 index] GTG ACT GGA GTT CAG ACG TGT GCT CTT CCG AT*C* T | HTO library |
| Library indexed reverse primer (Fragmented libraries) | *CAA GCA GAA GAC GGC ATA CGA GAT* [8bp i7 index] GTG ACT GGA GTT CAG ACG TGT | Regular mRNA library, long cDNA library |

**Supplementary Table 3:** The reference and annotation files used in the study.

| Type | File |
|---|---|
| Ensembl GTF | https://ftp.ensembl.org/pub/current_gtf/homo_sapiens/Homo_sapiens.GRCh38.109.gtf.gz |
| miRBase | https://mirbase.org/ftp/CURRENT/genomes/hsa.gff3 |
| Database of small human non-coding RNAs (DASHR) | https://dashr2.lisanwanglab.org/downloads/dashr.v2.sncRNA.annotation.hg38.gff |
| Enhancer RNA | https://fantom.gsc.riken.jp/5/datafiles/latest/extra/Enhancers/human_permissive_enhancers_phase_1_and_2.bed.gz |

**Supplementary Table 4:** Bioinformatics tools and packages used in the study.

| Tool/Package | Version | Application |
|---|---|---|
| cutadapt | 4.4 | Trimming and adapter removal |

| | | |
|---|---|---|
| riboDetector | 0.2.7 | rRNA sequences removal in silico |
| STARSolo | 2.7.10b | Alignment and single cell count matrix |
| Cite-seq-count | 1.4.0 | Demultiplexing HTO and ADT oligo barcode tags |
| RStudio | 4.3.1 | Seurat and downstream analysis of count matrix |
| future | 1.33.0 | Parallel threading to work with big single-cell matrix data |
| Seurat | 4.0 | Single-cell data analysis |

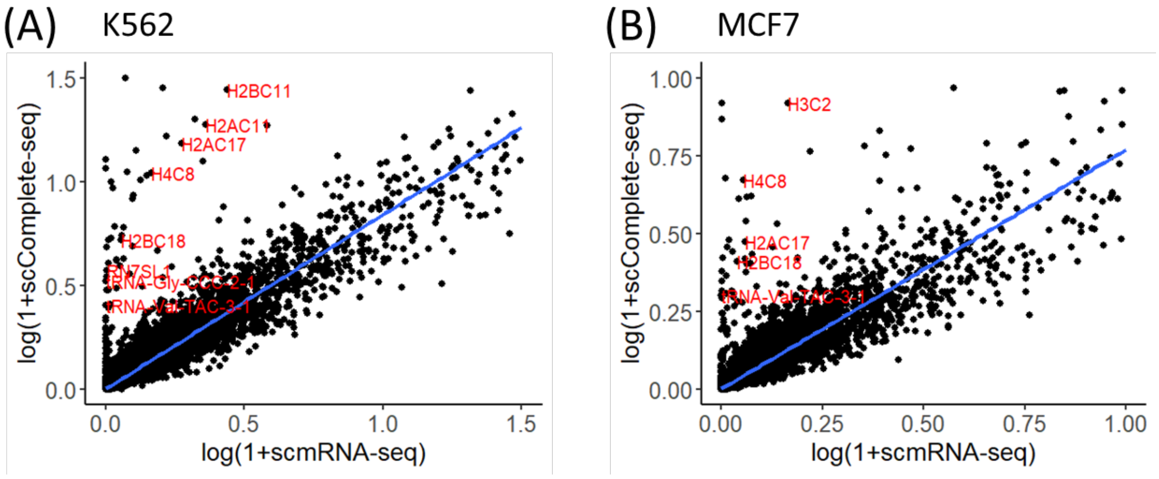


**Supplementary Figure 1:** Aggregated read counts obtained from scComplete-seq versus scmRNA-seq in (A) K562 and (B) MCF7 cells.

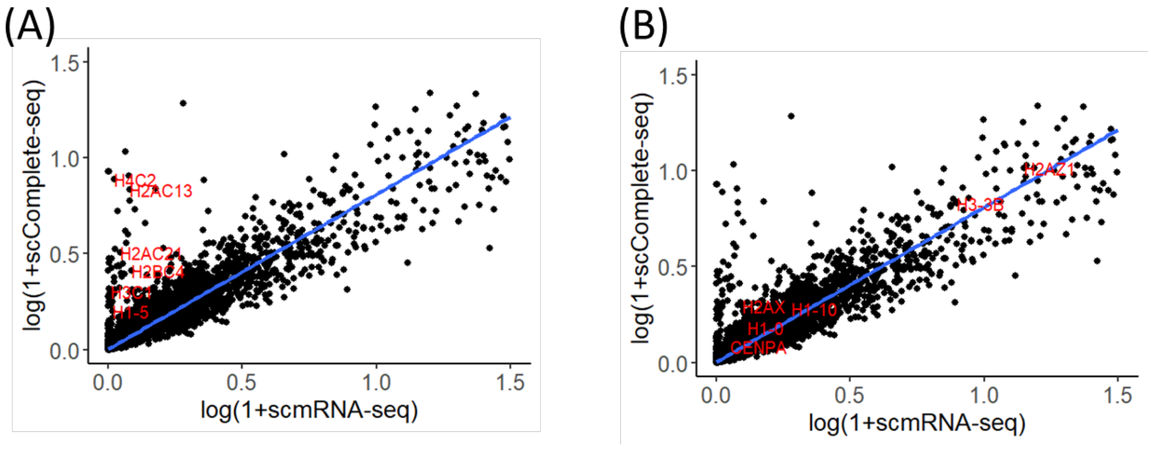


**Supplementary Figure 2:** Aggregated read counts obtained from scComplete-seq versus scmRNA-seq for HEK293T cells with (A) non-polyadenylated and (B) polyadenylated histone genes labeled.

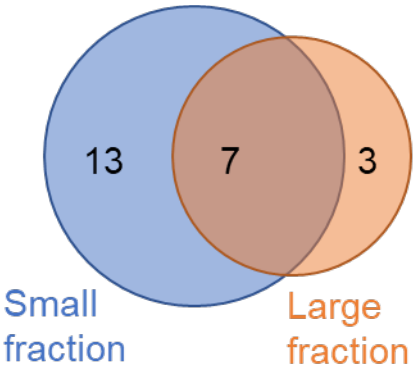


**Supplementary Figure 3:** Overlap between the short non-coding RNA in the top 50 DEG identified in the small and large fractions of scComplete-seq.

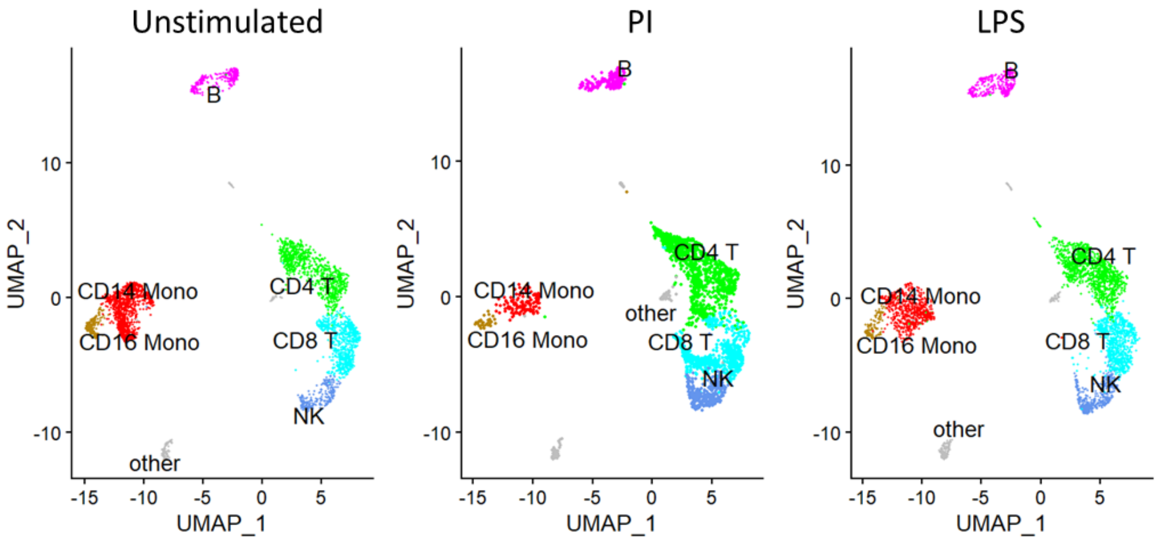


**Supplementary Figure 4:** UMAP after data integration showing the cell clusters obtained from unstimulated, PI-stimulated, and LPS-stimulated PBMC cells.

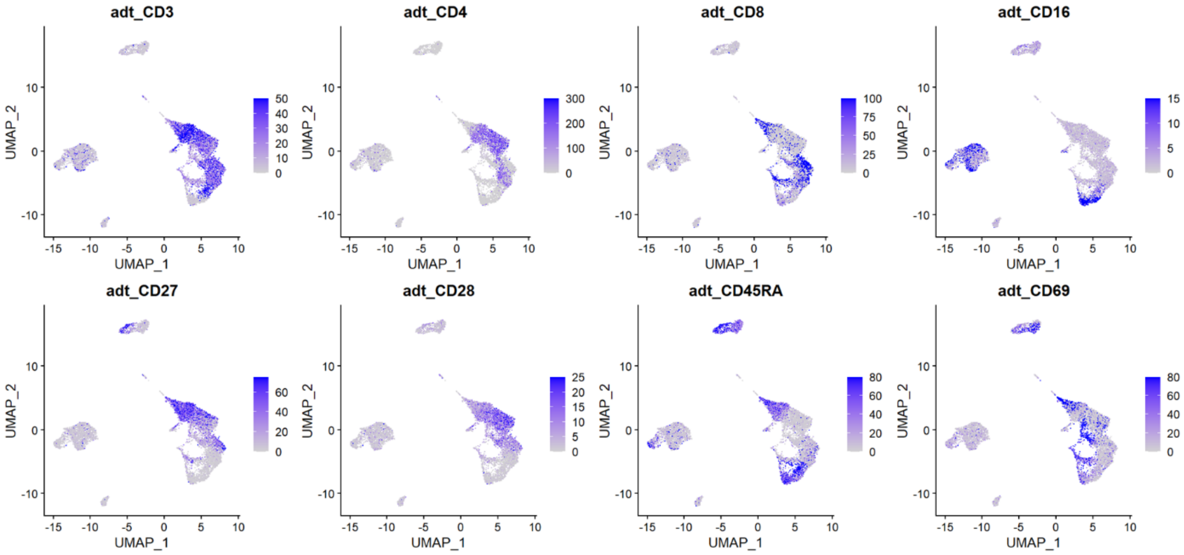


**Supplementary Figure 5**: Feature plots showing the antibody counts for the PBMC cells.

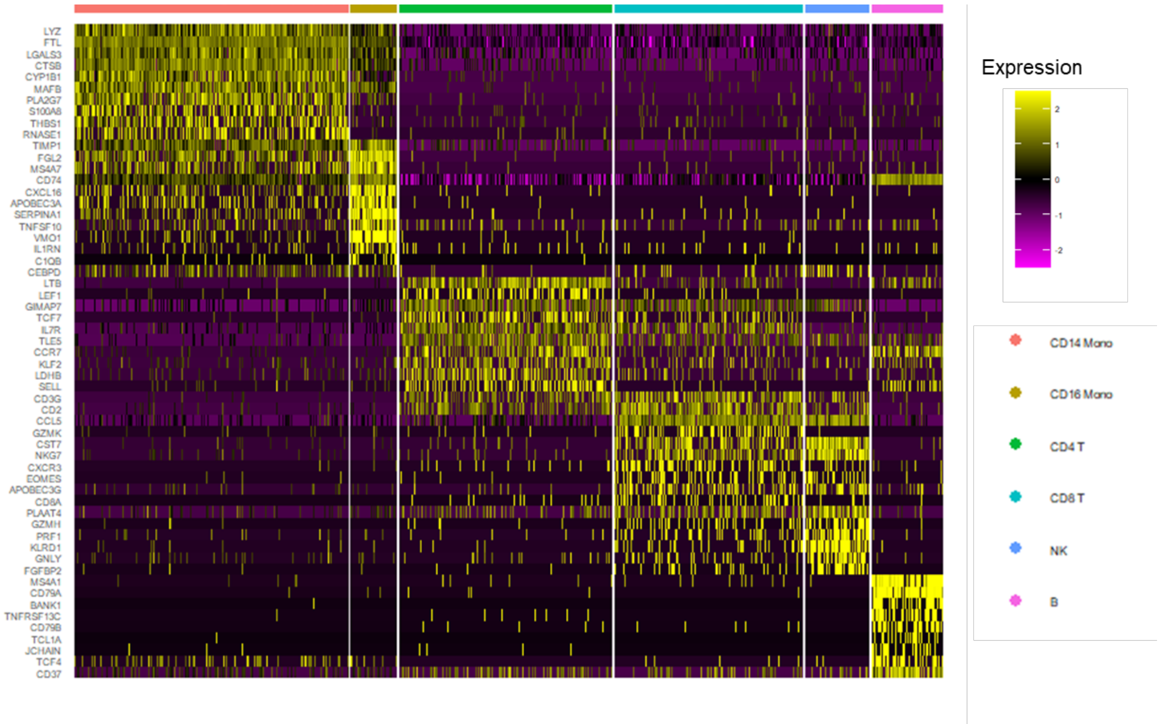


**Supplementary Figure 6**: Heatmap of protein-coding genes RNA expression obtained from the large fraction of scComplete-seq of PBMC cells.

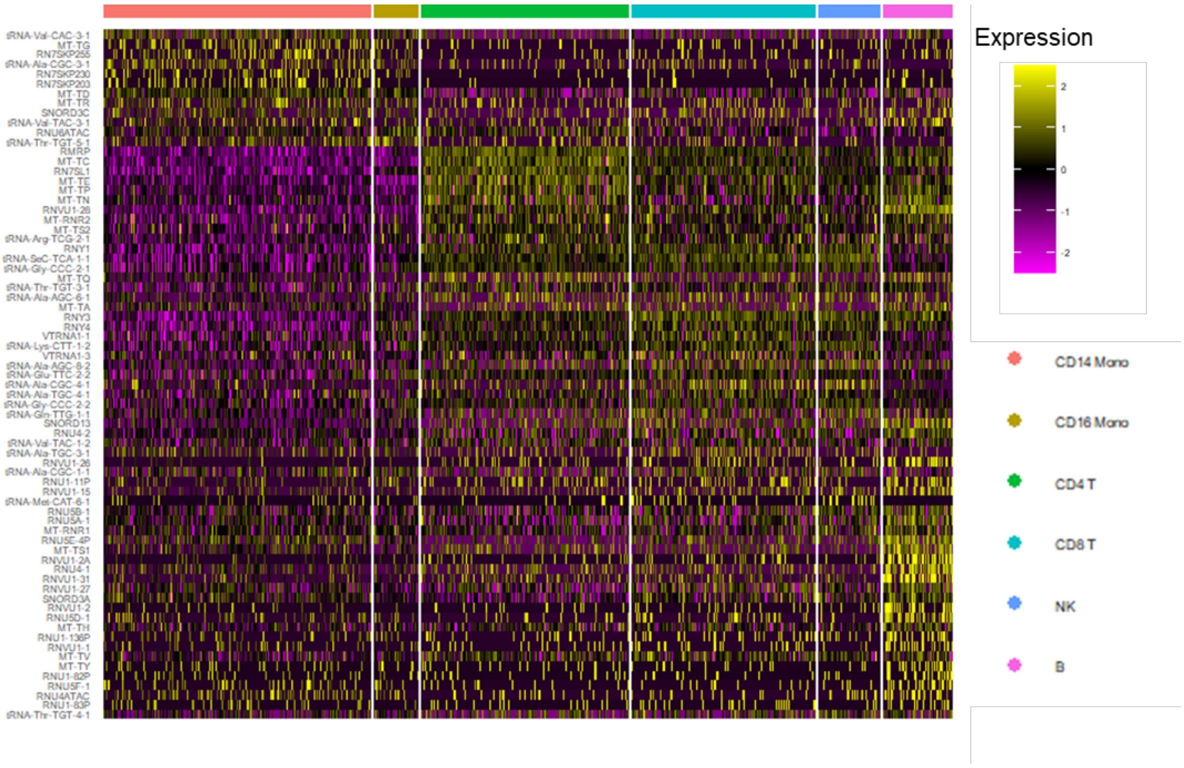


**Supplementary Figure 7:** Heatmap of short non-coding RNA expression obtained from the small fraction of scComplete-seq of PBMC cells.

**Adapter Trimming**

- Use cutadapt to trim polyA tail and GGG from TSO
- cutadapt -u 3 -o
- cutadapt -m 16 -a AAAAAAAAAAAAAAA

***in silico* rRNA removal**

- Use ribodetector to remove rRNA sequences
- ribodetector_cpu -l 147 -e rrna --chunk_size 256

**Alignment**

- Use STARSolo to perform alignment to custom annotation file and generate count matrix
- STAR-2.7.10b/bin/Linux_x86_64/STAR --runThreadN 16 --soloType CB_UMI_Simple --soloStrand Forward --outFilterMultimapNmax 50 --soloCBwhitelist /cellranger-7.1.0/lib/python/cellranger/barcodes/3M-february-2018.txt --soloUMIlen 12 --outFilterScoreMinOverLread 0 --outFilterMatchNminOverLread 0 --winAnchorMultimapNmax 100 --outFilterMatchNmin 15 --outFilterScoreMin 14 --outFilterMismatchNoverLmax 0.05 --seedMultimapNmax 100000 --seedSplitMin 9 --seedSearchStartLmax 25 --soloCellFilter EmptyDrops_CR --soloCBmatchWLtype 1MM_multi_Nbase_pseudocounts --soloUMIdedup 1MM_CR --soloFeatures Gene GeneFull SJ Velocyto --soloMultiMappers EM --limitBAMsortRAM 66243363923

**Hashtag and antibody feature couting**

- Use CITE-seq-Count to assign antibody and hashtag counts to respective cells
- CITE-seq-Count -cbf 1 -cbl 16 -umif 17 -umil 28 -t ADTtags/HTOtags.csv --bc_collapsing_dist 1 --umi_collapsing_dist 1 --max-error 1 -cells 16000 -wl GeneFullFilteredBarcodes.csv

**scComplete-seq data analysis**

- Use Seurat (v4.0)
- Separate large and small fraction cDNA as different assay

**Supplementary Figure 8**: Workflow for single-cell read data pre-processing, alignment, and raw count data retrieval and downstream processing. The important parameters of the tools were given in the commands.
